# A STAT3 Degrader Demonstrates Pre-clinical Efficacy in Venetoclax resistant Acute Myeloid Leukemia

**DOI:** 10.1101/2024.08.05.599788

**Authors:** Samarpana Chakraborty, Claudia Morganti, Bianca Rivera Pena, Hui Zhang, Divij Verma, Kimberly Zaldana, Nadege Gitego, Feiyang Ma, Srinivas Aluri, Kith Pradhan, Shanisha Gordon, Ioannis Mantzaris, Mendel Goldfinger, Eric Feldman, Kira Gritsman, Yang Shi, Stefan Hubner, Yi Hua Qiu, Brandon D. Brown, Anna Skwarska, Amit Verma, Marina Konopleva, Yoko Tabe, Evripidis Gavathiotis, Simona Colla, Jared Gollob, Joyoti Dey, Steven M Kornblau, Sergei B. Koralov, Keisuke Ito, Aditi Shastri

## Abstract

Acute myeloid leukemia (AML) is an aggressive hematologic malignancy that continues to have poor prognosis despite recent therapeutic advances. Venetoclax (Ven), a BCL2-inhibitor has shown a high response rate in AML; however, relapse is invariable due to mitochondrial dysregulation that includes upregulation of the antiapoptotic protein MCL1, a central mechanism of Ven resistance (Ven-res). We have previously demonstrated that the transcription factor STAT3 is upregulated in AML hematopoietic stem and progenitor cells (HSPCs) and can be effectively targeted to induce apoptosis of these aberrant cells. We now show that overexpression of STAT3 alone is sufficient to initiate a strong AML phenotype in a transgenic murine model. Phospho-proteomic data from Ven treated AML patients show a strong correlation of high total STAT3 and phospho-STAT3 [both p-STAT3(Y705) and p-STAT3(S727)] expression with worse survival and reduced remission duration. Additionally, significant upregulation of STAT3 was observed in Ven-res cell lines, in vivo models and primary patient samples. A novel and specific degrader of STAT3 demonstrated targeted reduction of total STAT3 and resulting inhibition of its active p-STAT3(Y705) and p-STAT3(S727) forms. Treatment with the STAT3 degrader induced apoptosis in parental and Ven-res AML cell lines and decreased mitochondrial depolarisation, and thereby dependency on MCL1 in Ven-res AML cell line, as observed by BH3 profiling assay. STAT3 degrader treatment also enhanced differentiation of myeloid and erythroid colonies in Ven-res peripheral blood mononuclear cells (PBMNCs). Upregulation of p-STAT3(S727) was also associated with pronounced mitochondrial structural and functional dysfunction in Ven-res cell lines, that were restored by STAT3 degradation. Treatment with a clinical-stage STAT3 degrader, KT-333 resulted in a significant reduction in STAT3 and MCL1 protein levels within two weeks of treatment in a cell derived xenograft model of Ven-res AML. Additionally, this treatment significant improvement in the survival of a Ven-res patient-derived xenograft *in-vivo* study. Degradation of STAT3 resulting in downregulation of MCL1 and improvements in global mitochondrial dysfunction suggests a novel mechanism of overcoming Ven-res in AML.

**Statement of Purpose:** Five-year survival from AML is dismal at 30%. Our prior research demonstrated STAT3 over-expression in AML HSPC’s to be associated with inferior survival. We now explore STAT3 over-expression in Ven-res AML, explain STAT3 mediated mitochondrial perturbations and describe a novel therapeutic strategy, STAT3 degradation to overcome Ven-res.

## Introduction

Acute Myeloid leukemia (AML) arises from malignant transformation of an immature hematopoietic precursor followed by clonal proliferation and accumulation of the transformed cells ^(1)^. The goal of therapy is to develop a treatment plan tailored to the molecular characteristics of the disease keeping in mind individual patient comorbidities and preferences. Ven is a selective inhibitor of the anti-apoptotic BCL2 protein. The FDA has approved Ven in combination with hypomethylating agents (HMAs) or low-dose cytarabine for treating newly diagnosed AML in patients over 75 years old or those ineligible for standard induction therapies. Ven is now also used extensively in the treatment of all frontline AML in combination with AML induction therapies such as Ven-FLAGIda, Ven 7+3 ^(2, 3)^. While these therapies are highly effective for inducing remissions, the 5-year survival in newly diagnosed AML remains dismal at 30%, with disease relapse being inevitable due to the outgrowth of therapy resistant hematopoietic stem and progenitor cells (HSPCs) ^(1)^. Relapse or refractory disease after exposure to Ven is a particularly difficult clinical problem to overcome and currently, there are no approved clinical approaches that have demonstrated efficacy after the failure of Ven.

Signal transducer and activator of transcription 3 (STAT3) belongs to the STAT family of transcription factors and is abberantly activated in several malignancies ^(4, 5)^. In response to proinflammatory cytokines such as IL-6 or growth factors, STAT3 undergoes phosphorylation and subsequent dimerization. Phosphorylation at Y705 leads to its translocation to the nucleus where it binds to STAT3 binding sequences within the promoter regions of target genes and is implicated in mediating oncogenic processes critical to cellular growth, proliferation and transformations etc. ^(6)^. Additionally, a pool of STAT3 is also present in the mitochondria, known as mitoSTAT3, that regulates activity of electron transport chain (ETC), and cellular respiration. In contrast to its role in nucleus, mitoSTAT3 undergoes phosphorylation at S727 residue and is associated with malignant transformations in prostate cancer, chronic lymphocytic leukemia, and breast cancer ^(7)^. However, the mitochondrial role of p-STAT3(Y705) is poorly understood in AML. While STAT3 has been an attractive target of research for therapeutic development for decades, the lack of specificity has led to off target effects and poor bioavailability of previously developed STAT3 inhibitors leading to their failure in clinical trials ^(8)^.

We have previously demonstrated that STAT3 is frequently de-methylated and overexpressed in myelodysplastic syndrome (MDS) & AML HSPCs and is associated with an adverse prognosis^(1)^. We have also reported that STAT3 transcriptionally controls several important leukemic drivers that emerge during Ven-res, such as the anti-apoptotic protein myeloid cell leukemia-1 (MCL1). MCL1 overexpression is the central mechanism of resistance to BCL2 inhibition in AML. MCL1 is a well-known direct transcriptional target of STAT3 and it is likely that changes to STAT3 will affect downstream MCL1 activity ^(9)^.

We now present data demonstrating that hyperactivation of STAT3 alone is sufficient to induce myeloid bias and promote leukemia in a transgenic murine model. We have observed worse overall survival and reduced remission duration when total STAT3 and its phosphorylated forms are upregulated in AML patients treated with Ven. Additionally, we show that both total and phosphorylated forms of STAT3 are upregulated in Ven-res AML. Notably, the serine phosphorylated form of STAT3 is associated with mitochondrial dysfunction in Ven-res AML. Importantly, a novel and clinically relevant STAT3 degrader effectively reverses the Ven-res phenotype in AML cells, improving survival in Ven-res PDX model and mitigating aberrant mitochondrial function in leukemic HSPCs.

## Results

### STAT3 overexpression drives myeloid bias

To understand the role of STAT3 in initiation and progression of leukemia, we have developed a transgenic murine model with a hematopoietic-specific hyperactivation of the STAT3 pathway (STAT3C-vavCre model). The murine model was generated by crossing R26STAT3Cstop^fl/fl^ mice with vavCre transgenic mice ^(10)^. STAT3C-vavCre double transgenic mice (n=17) were validated by GFP expression in HSPCs and differentiated hematopoietic cells. The STAT3C-vavCre mice developed ruffled fur, a hunched phenotype and weight-loss by 24-26 weeks of age ^(11)^. Histopathologic analysis of the STAT3C-vavCre mice revealed complete destruction of splenic architecture with a spleen engorged with dysplastic appearing leukemic blasts and significant splenomegaly and hepatomegaly as compared to compared to STAT3-only and VavCre-only controls (Figure 1 A-C). Complete Blood Count (CBC) analysis of STAT3C-vavCre mice showed significant anemia and progression to a proliferative phenotype with higher white blood cells (WBC) (Figure 1 D-F). Using hematopoietic stem cell (HSC) lineage analysis by FACS on the BM of the mice, we observed that STAT3C-vavCre mice drive the selective expansion of multipotent progenitor population (MPP) and long-term hematopoietic stem cell (LT-HSC) population compared to controls, suggesting a stem and progenitor phenotype, as seen in high risk MDS/AML (Figure 1 G). Serial transplantation experiment using BM cells from STAT3C-VavCre donor mice in irradiated C57BL/6 recipient mice showed significant transplantation ability, demonstrating the proliferative capacity of STAT3 overexpression mice (Figure 1H, I). Increased percentage of myeloid and neutrophil cells was observed in STAT3C-vavCre mice as compared to WT mice using ExCITE-Seq, suggesting a strong and sustained myeloid bias that is typically observed in AML (Figure 1 J-M).

**Figure 1:**
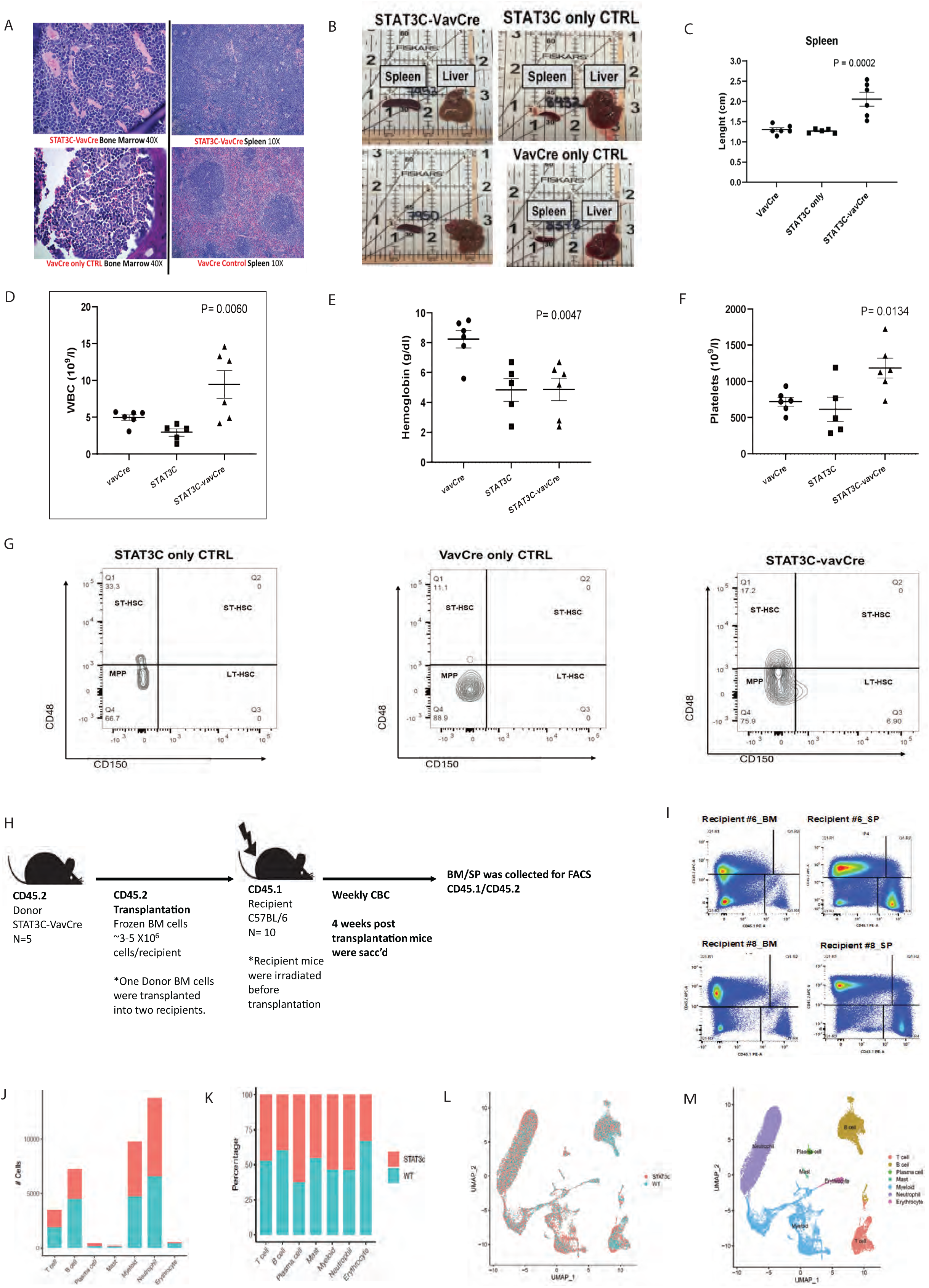
STAT3 overexpression is sufficient to develop leukemia phenotype and myeloid bias in murine model. A) Histopathologic image of the STAT3C-vavCre mice vs control shows complete destruction of splenic architecture with leukemic blasts. B, C) Significant splenomegaly and hepatomegaly is observed in in STAT3C-vavCre mice. D-F) Complete Blood Count (CBC) analysis of STAT3C-vavCre mice showed increased WBC count, significant anemia and thrombocythemia, as compared to control mice. G) FACS HSC Lineage analysis on the BM of STAT3C-vavCre mice shows the selective expansion of myeloid progenitor population (MPP) and long term hematopoietic stem cell (LT-HSC) population compared to controls. H) Competitive transplantation model was established using BM cells from STAT3C-VavCre donor mice transplanted in recipient C57BL/6 mice. I) FACS of CD45.1 vs CD45.2 markers demonstrates high transplantation ability of STAT3Cvav-Cre mice in two recipient C57BL/6 mice (#6 and #8). J) ECCITE seq data shows more number of myeloid and neutrophil cells as compared to other cell types in STAT3C-vavCre mice. K) STAT3C-vavCre mice also shows higher percentage of myeloid and neutrophil cells in STAT3C-vavCre mice as compared to WT mice, suggesting myeloid bias that is typically observed in AML. L, M) UMAP showing enriched clusters specific to myeloid and neutrophil cell types in STAT3C-vavCre mice as compared to WT mice.

### STAT3 is overexpressed in Venetoclax resistant AML

To understand the implication of Ven-res *in-vitro*, we developed Ven-res AML cell lines – MOLM13, MV411 and lymphoma cell line SU-DHL1 by treating the parental cells with increasing dose of Ven over a period of 6-8 weeks. In the Ven-res cell lines, we observed increased total STAT3 along with one of its downstream effector, MCL1, when compared to parental cell lines (Figure A). qPCR validated our findings and additionally demonstrated increased expression of BCL2 and BCL-xl in MOLM13 Ven-res cells (Figure 2B). Furthermore, clinical data from AML patients that received treatment with Ven containing regimens has a significantly worse outcome if they had high expression of total STAT3 along with increased phosphorylation for p-STAT3(Y705) and p-STAT3(S727) for both overall survival (OS, n=138 Y705 p = 0.01, S727 p=0.0005) and remission duration (RemDur, n=90, p<0.05) (Figure 2 C, D; Supplementary Figure 1 A, B). BM sample from AML patients shows significantly higher expression of STAT3 (p=0.023) in specific clusters that emerge post Ven and decitabine treatment in therapy-resistance/relapse as observed by bulk RNAseq (Figure 2E, Table 1).

**Figure 2:**
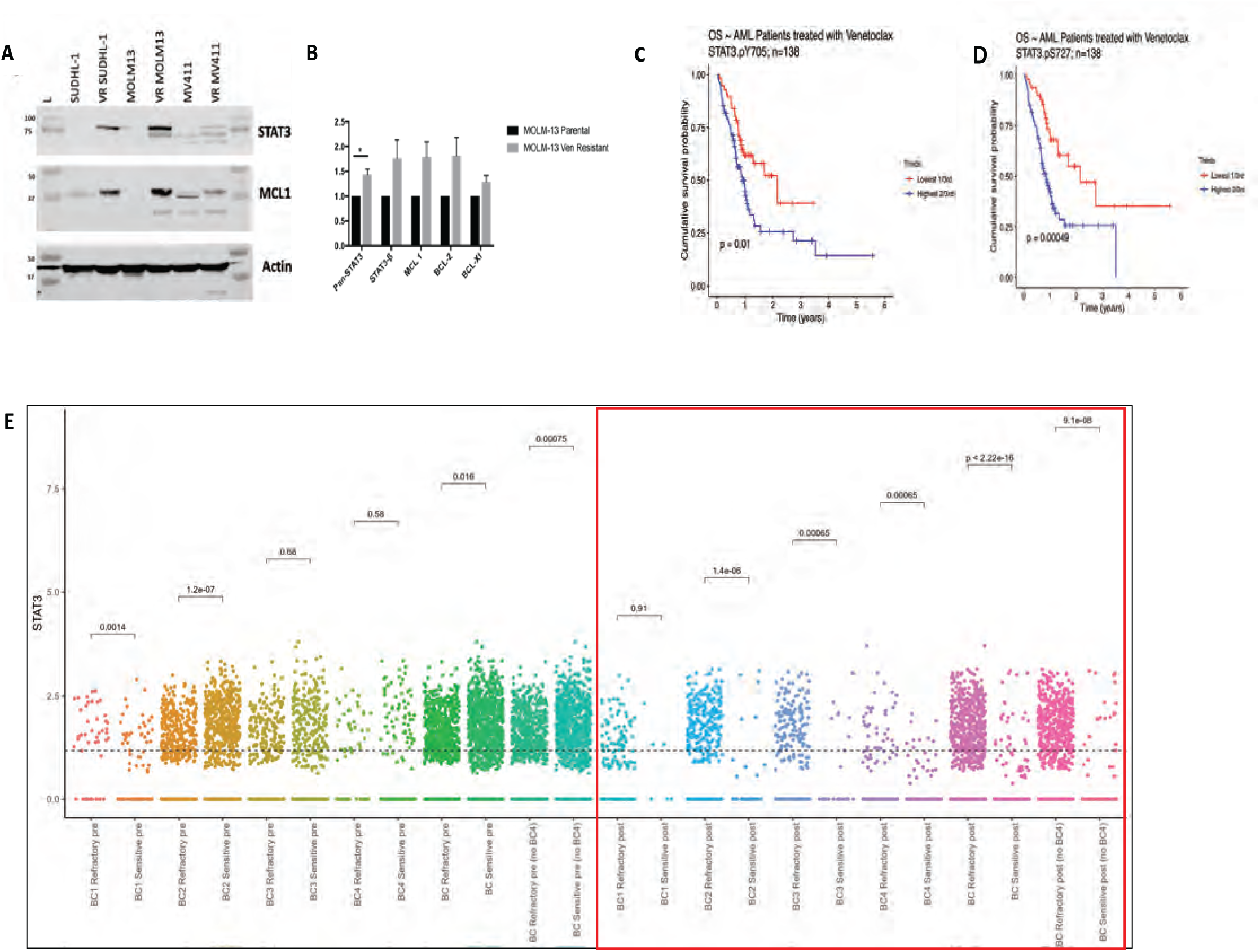
STAT3 is overexpressed in Venetoclax resistant AML. A) Increased expression of total STAT3, p-STAT3(Y705) and MCL1 was observed in Ven resistant MOLM13, MV411 and SU-DHL1 cell lines as compared to their parental cell lines. B) RT-PCR using RNA from parental and Ven-res MOLM13 cell lines shows upregulation of total STAT3, STAT3-β, MCL1, BCL2 and BCL-xl in MOLM13 Ven-res cells. C, D) Phospho-proteomic analysis on AML patients treated with Ven shows significant worse OS with higher expression of p-STAT3(Y705) and p-STAT3(S727), respectively. E) Bulk RNA seq data on BM samples from post Ven/decitabine treated relapsed/refractory AML patients shows significant higher expression of STAT3.

**Tables 1:** STAT3 gene expression at DEC/VEN timing; non-responder (n=3) vs initial responder (n=7) in relapsed cases.

| Average of Gene expression | Non-responder | responder | p value |
| --- | --- | --- | --- |
| Stat3 | 9932.219 | 7968.585 | 0.200 |
| Stat3 / ABL1 (corrected by ABL1) | 3.539 | 2.136 | **0.023** |

### STAT3 degraders effectively and selectively degrade STAT3, initiate apoptosis and reduce dependency on MCL1 for survival in Ven-res AML

As our data shows high STAT3 expression in Ven-res AML, we hypothesized that STAT3 degradation can be a novel therapeutic strategy against Ven-res AML. To this effect, we tested highly specific potent heterobifunctional tool degraders of STAT3 ^(16,17)^ (KTX-201, KTX-105). The selective degradation of STAT3 in both parental and Ven-res MOLM13 cell line was confirmed using western blot on treatment with two STAT3 degraders (D1: KTX-201, D2: KTX-105) at 0.1, 1 and 10μM doses for 24 hours, with no effect on STAT3 levels when treated with structural controls (C1, C2) or DMSO (Figure 3 A, B). Additionally, we confirmed these compounds selectively targeted STAT3 but not STAT5 (Supplementary Figure 2 A, B), suggesting target specificity. Treatment with KTX-201 resulted in a dose dependent decrease in proliferation of cell lines that express high levels of STAT3 such as SU-DHL1 (lymphoma cell line) as well as MOLM16 (AML cell line) at nanomolar concentrations (Figure 3C, D). KTX-201 treatment also led to significant induction of apoptosis in both MOLM13 parental and Ven-res AML cell lines at 48 hours (Figure 3E). BH3 profiling of Ven-res MOLM-13 supports an increased dependency on MCL1 as observed by increase in mitochondrial depolarization on treatment with MS1 and NOXA peptides, which bind specifically to MCL1 (Figure 3F). Interestingly, we observe that treatment with KTX-201 leads to approximately 20% reduction in the mitochondrial depolarization of MS1 and NOXA, thus reducing the dominion of MCL1 in mediating Ven-res (Figure 3G).

**Figure 3.**
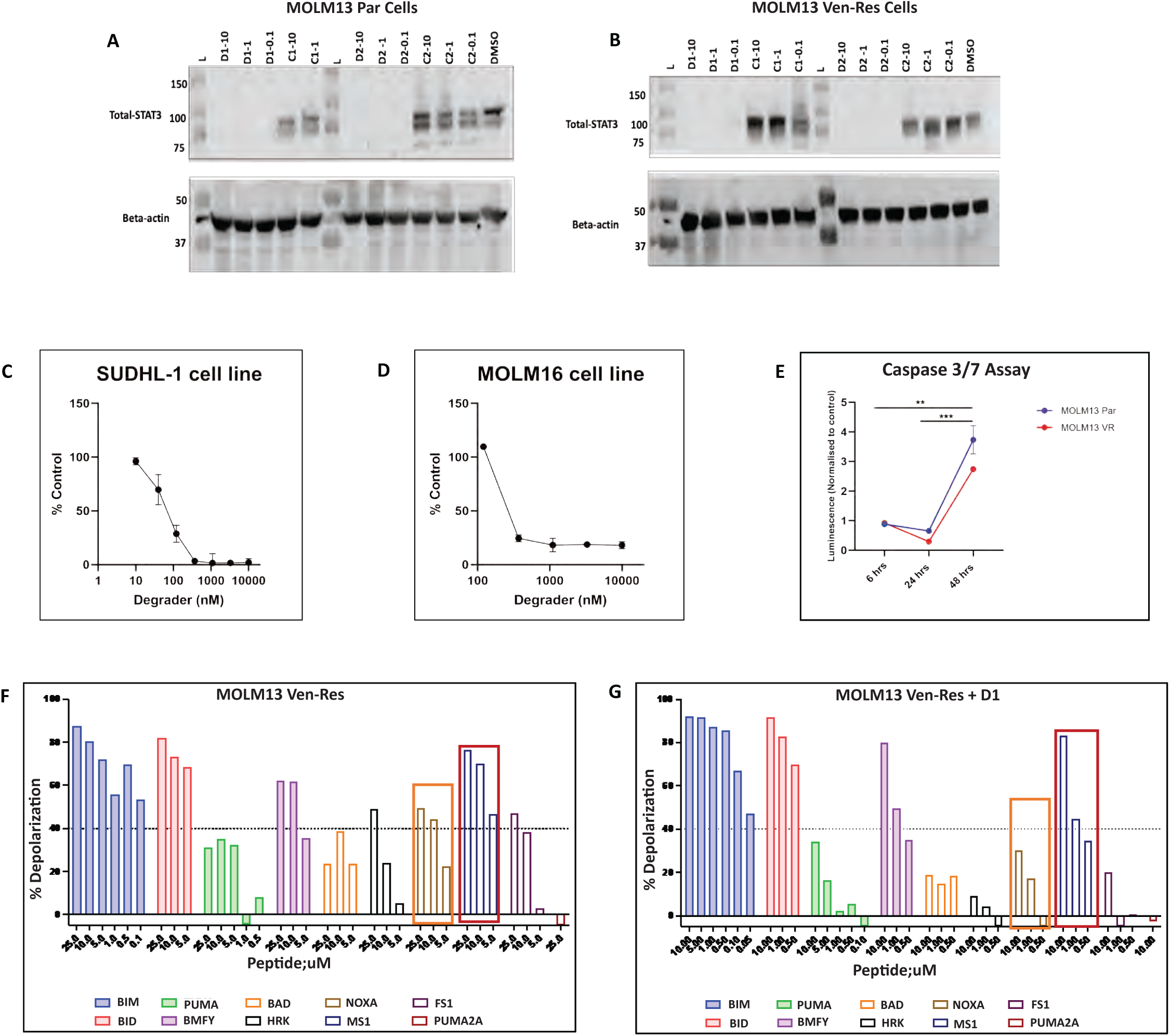
Selective degradation of STAT3 by degraders occur via initiating apoptosis and reduces dependency on MCL1 in Ven-res AML. A, B) Western blots showing selective degradation of STAT3 in both parental and Ven-res MOLM13 cell line on treatment with STAT3 degraders (D1: KTX-201, D2: KTX-105) at 0.1, 1 and 10μM doses for 24 hours, with no effect on STAT3 levels when treated with structural controls (C1, C2) or DMSO. C, D) Dose dependent decrease in proliferation of SU-DHL1 and MOLM-16 cell lines 72 hours post treatment with nanomolar concentrations of KTX-201, measured by CTG assay. E) KTX-201 treatment led to significant induction of apoptosis in both MOLM13 parental and Ven-res AML cell lines at 48 hours, as measured by Caspase 3/7 assay. F) BH3 profiling of Ven-res MOLM-13 shows increased depolarization when treated with MS1 and NOXA peptides even at 5μM concentration, suggesting an increased dependency on MCL1 protein. G) Treatment with KTX-201 led to reduction in the MS1 and NOXA induced mitochondrial depolarization in MOLM13-Ven-res cell line.

### STAT3 degrader KTX-201 is taken up by primary AML and Ven res AML PBMNCs and leads to STAT3 downregulation and effective erythroid and myeloid differentiation

In primary Ven-res AML patient samples, we have noted effective degradation of STAT3 (>90%) in multiple patient PBMNC samples on treatment (D1: KTX-201, D2: KTX-105) for 24 hours (Figure 4A-C), with no change in STAT3 levels when treated with structural controls (C1, C2) or DMSO.

**Figure 4.**
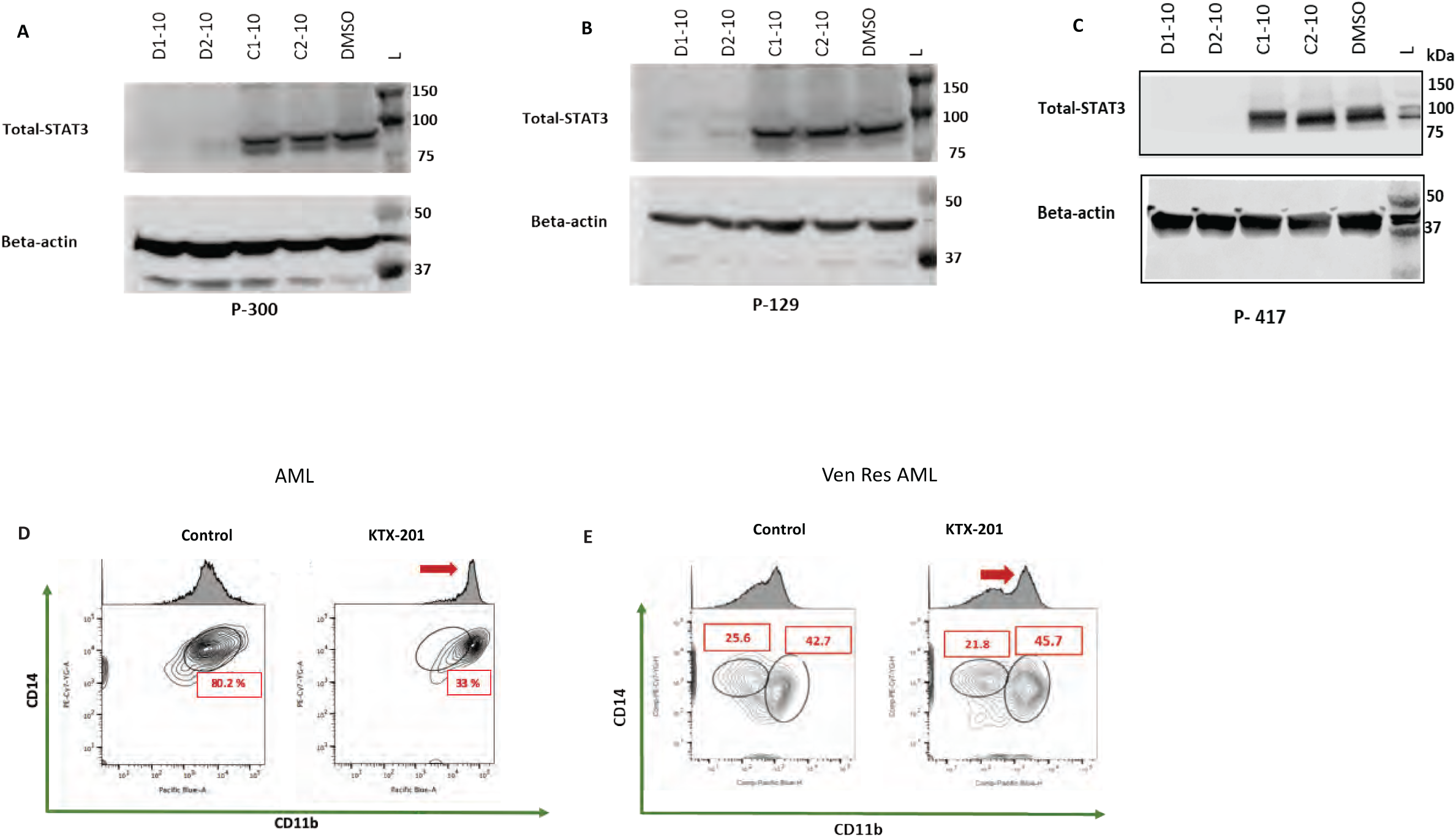
Primary AML and Ven-res AML PBMNCS show effective degradation of STAT3 on treatment with KTX-201 and induce erythroid and myeloid differentiation. A, B) Western blots showing effective degradation of total STAT3 in AML patient PBMNC and C) Ven-res AML patient PBMNC samples on treatment with STAT3 degraders - D1: KTX-201, D2: KTX-105 for 24 hours, with no change in STAT3 levels when treated with structural controls (C1, C2) or DMSO. D) FACS post CFU assay using AML patient PBMNC shows increased differentiation of myeloid markers CD11b and CD14, when treated with KTX-201 as indicated by red arrow. E) FACS post CFU assay in Ven-res AML patient PBMNC shows increased differentiation of myeloid markers CD11b and CD14, when treated with KTX-201 as indicated by red arrow.

Stem and progenitor cells of MDS and AML show arrested and dysplastic differentiation ^(18, 19)^. We treated a number of primary AML and Ven-res AML patient PBMNCs samples with KTX-201 and control and grew them in methylcellulose assays supplemented with cytokines. Cells were harvested after 14 days and assessed for erythroid and myeloid differentiation by flow cytometry. We observed increased erythroid (∼ 1.5 fold) and myeloid differentiation (∼ 2.5 fold) on treatment with KTX-201 as measured by CD71, Glycophorin A that are markers of early erythroid differentiation and CD14, CD11b that are myeloid differentiation markers, especially in Ven-res AML patients (Figure 4D, E; Supplementary Figure 3A, B). Interestingly, no differentiation was observed in healthy samples (Supplementary Figure 3C), suggesting the specificity of the degrader to AML stem and progenitor cells with no effect on normal cell types.

### Venetoclax resistance increases p-STAT3(S727) protein levels in vitro and leads to structural alterations in mitochondria

Previous report by Chen et al. have suggested a possible structural and functional changes in mitochondria on Ven treatment ^(20)^. The antiapoptotic protein MCL1, known to localize in the mitochondria, is upregulated and plays a key role in the development of mitochondrially mediated Ven-res ^(9,14,21)^. Additionally, literature also suggests presence of STAT3 in the mitochondria (mitoSTAT3) where it undergoes phosphorylation at Ser727 residue; p-STAT3 (S727) ^(22)^. MitoSTAT3 is known to be important for maintaining optimal mitochondrial respiration by modulating complex I and II of the ETC ^(22)^ and prevents ROS generated oxidative stress in the heart ^(23,24)^. It has been studied extensively in cancers such as chronic lymphoid leukemia, pancreatic and breast cancers ^(25–27)^, in addition to cardiovascular ^(28, 29)^ and neuronal research ^(30)^. We hypothesized that in addition to MCL1, p-STAT3(S727) is also upregulated and causes mitochondrial dysregulation in Ven-res AML. To confirm our hypothesis, we performed western blot on MOLM13 parental and Ven-res cells and probed for p-STAT3(S727) levels.

Ven-res MOLM13 cells show an increased expression of p-STAT3(S727) on western blot as compared to MOLM13 parental cells (Figure 5A). On comparing the ultrastructure of mitochondria using electron microscopy (EM), we observed swollen node-like structure of the cristae of MOLM13 Ven-res cells, that were absent in the MOLM13 parental cells (Figure 5B). Cristae are present in the inner membrane of the mitochondria where oxidative phosphorylation (OXPHOS)/ATP synthesis occurs and therefore, abnormalities in the cristae structure affect mitochondrial bioenergetics ^(31, 32)^. Quantification of mitochondrial structure suggests an increased cristae total area as compared to the Ven-res cell lines (Figure 5C). We further checked if STAT3 localizes in the mitochondria using immunofluorescence, and observed a significant increase in the intensity of both total STAT3 and p-STAT3(S727) in the mitochondria (Figure 5 D-I). To validate our findings of mitochondrial structural abnormalities in primary patient samples, we collected fresh PBMNCs from Ven-res patient samples and healthy subjects, fixed the cells and subjected them to EM imaging, followed by image analysis (Figure 5J). The significant increase in cristae area, width along with cristae total area compared to mitochondrial area in the Ven-res sample confirm our *in-vitro* findings in primary patient samples (Figure 5 K-N).

**Figure 5.**
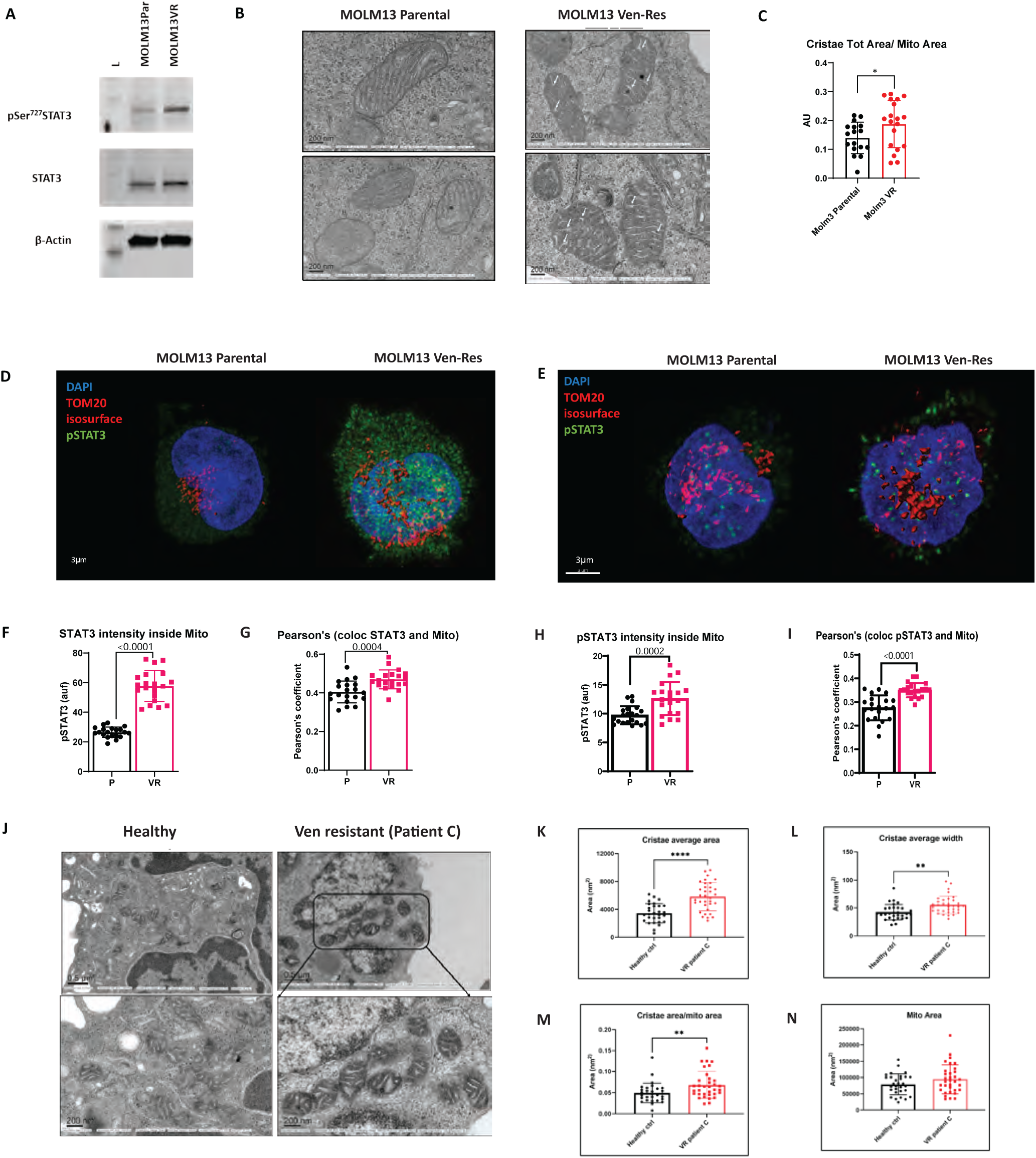
Overexpression of p-STAT3(S727)and mitochondrial structural alterations in Ven-res AML. A) Western blot on MOLM13 parental and Ven-res cells shows increased expression of p-STAT3(S727) levels in Ven-res. B) EM imaging of MOLM13 Ven-res cell lines show swollen node like structure in the mitochondria cristae of Ven-res cells, as compared to MOLM13 parental cells. C) Quantification of the EM image shows significant increase in cristae total area as compared to mitochondrial area in the MOLM13 Ven-res cell line, as compared to MOLM13 parental cells. D, E) Immunofluorescence image shows increased colocalization of total STAT3 and p-STAT3(S727) with TOM20, in the mitochondria of MOLM13 Ven-res cell line. TOM20 is a protein that belongs to the import receptor complex of the mitochondrial outer membrane. F-I) Quantification of the IF signals shows a significant increase in the intensity of both total STAT3 and p-STAT3(S727) in the mitochondria with significant Pearson’s co-localization coefficient. J) EM image of PBMNCs from Ven-res patient (patient C) shows huge nodule like cristae in mitochondria as compared to healthy subject. K-N) Image analysis shows a significant increase in cristae area, width along with cristae total area as compared to mitochondrial area in the Ven-res patient PBMNC sample as compared to healthy subject (Figure 5 K-N).

### Treatment with STAT3 degrader can reduce total and mitoSTAT3 inside mitochondria and reverse mitochondrial dysfunction in Ven-res

After establishing that p-STAT3(S727) is upregulated in Ven-res, we wanted to check if a STAT3 degrader can mitigate the damage caused by elevated mitoSTAT3. We observed that expression of p-STAT3(S727) undergoes significant reduction in MOLM13 Ven-res cells as compared to MOLM13 parental cells on treatment with 100nM D1 (KTX-201) (Fig 6A). We hypothesize that on treatment with the STAT3 degrader, a global reduction in total STAT3 also greatly reduces the phosphorylated active forms of STAT3 in both mitochondria p-STAT3(S727) as well as cytoplasm p-STAT3(Y705). The resulting loss of STAT3 signaling are likely responsible for structural and functional changes in mitochondria. To understand if STAT3 degrader can reverse mitochondrial dysfunction in vitro, we performed subcellular fractionation using density gradient ultracentrifugation to isolate mitochondria in MOLM13 Ven-res cell line, with and without treatment with KTX-201. Western blot of the protein isolates showed high expression of total STAT3 as well as p-STAT3(S727) in the mitochondria of the untreated cells, that undergo significant decrease on treatment with KTX-201 (Figure 6B). Densitometry of the blot showed downregulation of STAT3, p-STAT3(S727) and MCL1 on treatment with KTX-201 (Figure 6C). Additionally, we isolated fresh PBMNCs from Ven-res patients and fixed the cells pre and post treatment with KTX-201 and performed EM imaging (Figure 6D). We observed long mitochondria in the untreated sample suggesting fission/fusion defects in the mitochondria, which post treatment with degrader leads to round/smaller fragments suggesting possible priming of the cells towards apoptosis. Treatment with KTX-201 also reduced cristae count, cristae area and cristae width per mitochondria (Figure 6E). Interestingly, image quantitation demonstrated that the decrease in mitochondrial area post treatment was comparable to that of healthy controls suggesting that STAT3 degradation can reverse the structural defects in mitochondria caused due to Ven-res (Figure 6F).

**Figure 6.**
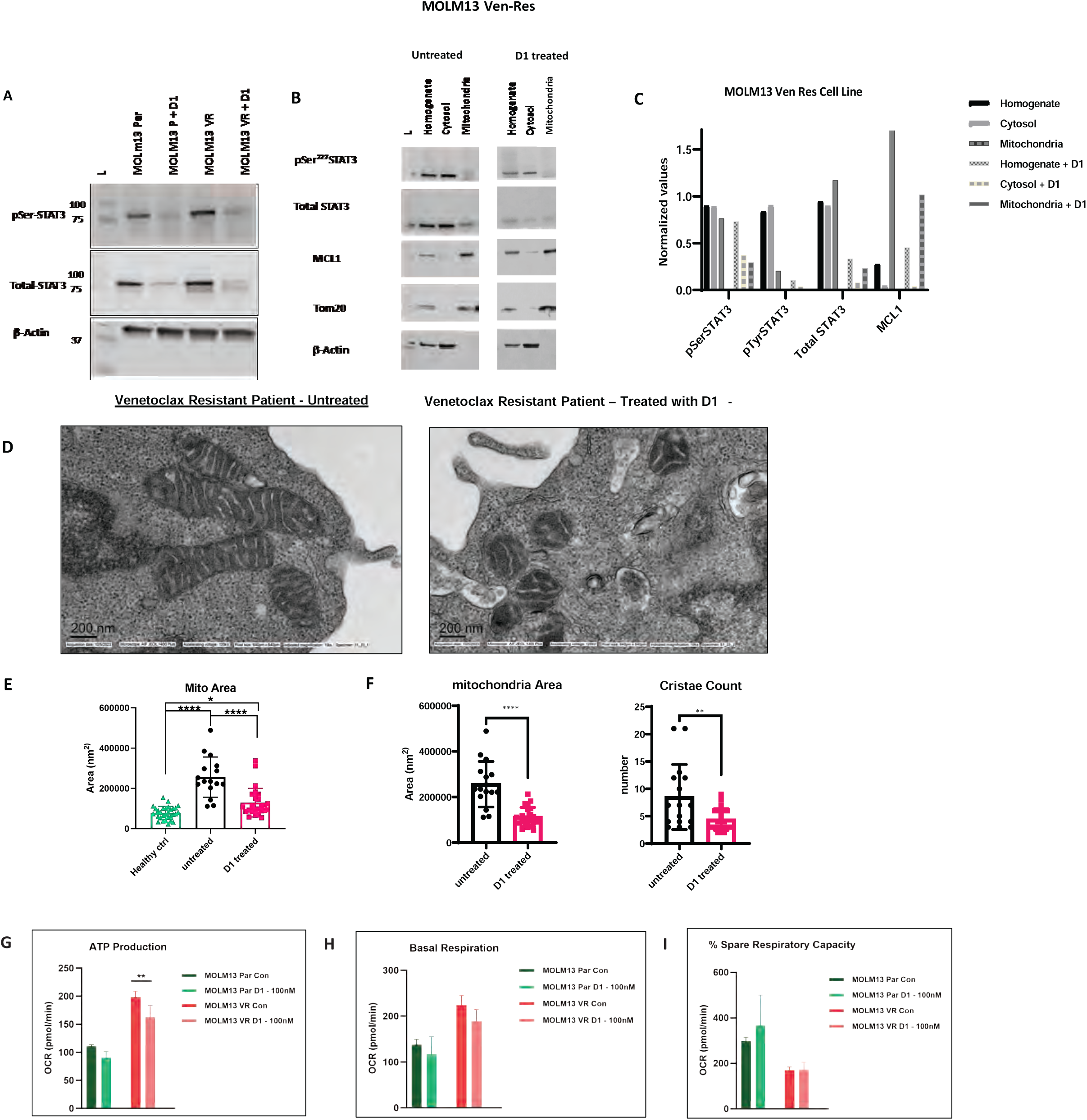
STAT3 degrader treatment can decrease mitoSTAT3 and mitigate mitochondrial dysfunction in venetoclax resistance. A) Western blot showing significant reduction in the expression of p-STAT3(S727) in MOLM13 Ven-res cells as compared to MOLM13 parental cells on treatment with 100nM D1 (KTX-201) for 24 hours. B) Western blot on subcellular fractionated protein isolates from MOLM13 Ven-res cell line, pre and post treatment with KTX-201, shows high expression of total STAT3 as well as p-STAT3(S727) in the mitochondria of the untreated cells, that undergo significant decrease on treatment with KTX-201. C) Densitometry of the western blot show downregulation of STAT3, p-STAT3(S727) and MCL1, especially in the mitochondrial fraction post treatment with KTX-201. D) EM imaging on PBMNCs from Ven-res patient pre and post treatment with KTX-201 shows altered structure of mitochondria in the untreated sample suggesting fission/fusion defect, which reverses post treatment with KTX-201. E) Image quantification of panel D shows significant decrease in mitochondrial area in Ven-res sample post treatment with KTX-201, and is comparable to that of healthy control. F) Treatment with KTX-201 also significantly reduced mitochondria area, cristae count, cristae area and cristae width per mitochondria in Ven-res PBMNC treated with KTX-201. G, H) Mito stress test using MOLM13 parental and Ven-res cells, pre and post treatment with D1 (KTX-201) shows an increase in ATP production and basal respiration in MOLM13 Ven-res cells as compared to parental cells. Treatment with KTX-201 caused a significant decrease in ATP production and basal respiration in MOLM13 Ven-res cells I) Mito stress test using MOLM13 parental and Ven-res cells, pre and post treatment with D1 (KTX-201) shows decrease in % spare respiratory capacity of MOLM13 Ven-res cells as compared to parental cell line. Minor change in % spare respiratory capacity was observed post STAT3 degrader treatment.

To decipher the functional consequence of mitochondrial structural defect in Ven-res, we performed functional analysis using Seahorse assay for MOLM13 parental and Ven-res cells, pre and post treatment with D1 (KTX-201). MOLM13 Ven-res cells showed a higher ATP production and basal respiration suggesting they have higher metabolic activity as compared to parental cells ^(33)^. This correlates with the concurrent increase in the cristae area of Ven-res cells observed that signify an increase in mitochondrial respiration. However, lower spare respiratory capacity of the resistant cells as compared to the parental cells suggests their declined ability to tolerate stress ^(34)^. Treatment with KTX-201 caused a significant decrease in ATP production and basal respiration in MOLM13 Ven-res cells (Figure 6G-I).

### Significant increase in survival was observed on treatment with STAT3 degrader in PDX model of Venetoclax resistance

We evaluated the pharmacodynamic activity of KT-333, a clinical stage STAT3 degrader (currently in Phase 1 clinical trial: NCT05225584) in MOLM13 Ven-res, a cell-derived xenograft (CDX) model of systemic Ven-res AML (Figure 7A). Spleens from these mice showed significant reduction of p-STAT3(Y705) (∼ 60%), total STAT3 (>90%) as well as MCL1 (∼ 70%), on treatment with two doses of KT-333, as assessed in week 2 post-transplant, 48 hours post drug administration (Figure 7B, C).

**Figure 7.**
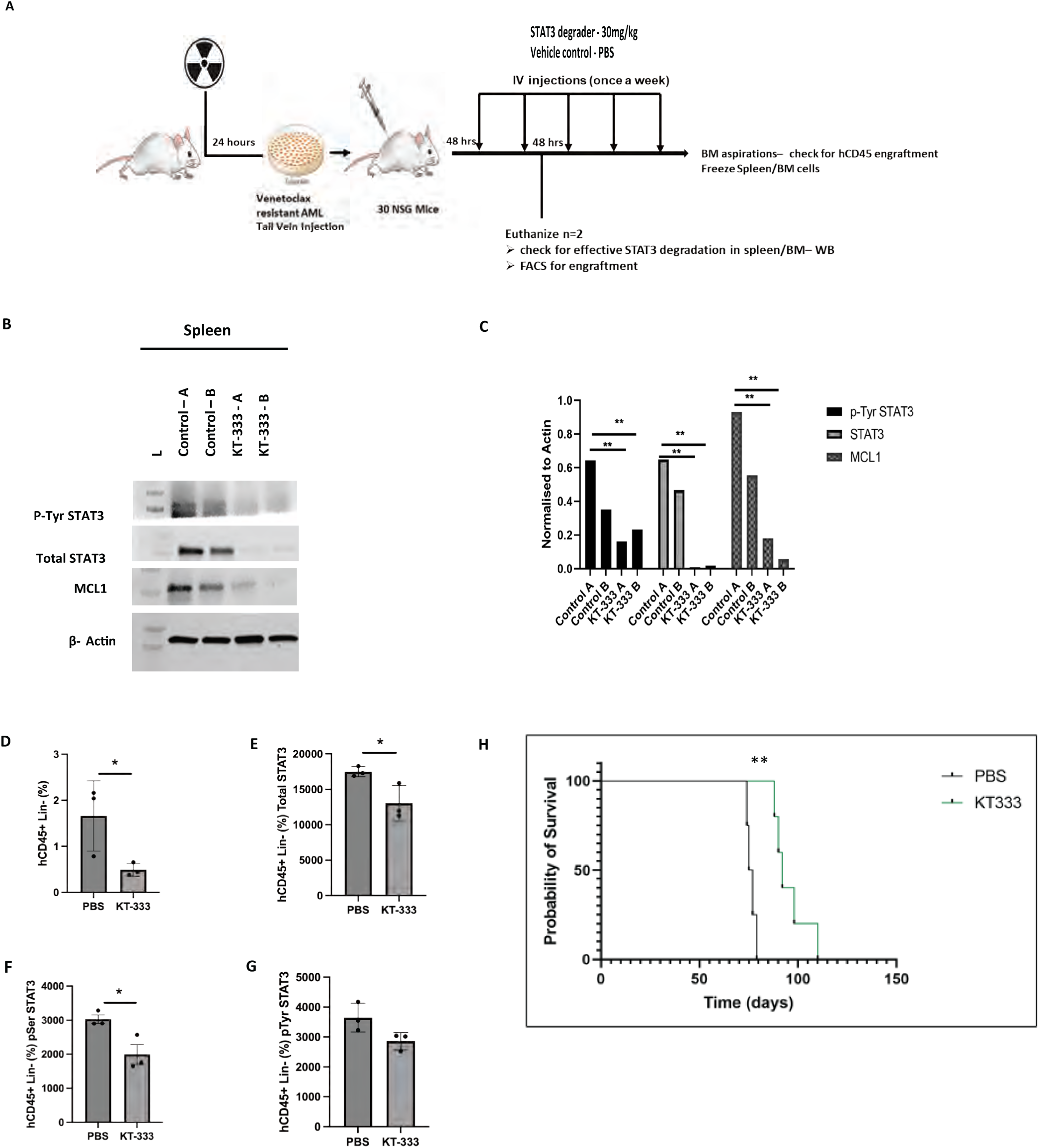
CDX and PDX model of venetoclax resistance show significant impact of STAT3 degrader in Ven-res AML. A) Schematic showing the development of cell-derived xenograft (CDX) and patient derived xenograft model of systemic Ven-res AML. B, C) Western blot and its densitometry showing significant reduction of p-STAT3(Y705), total STAT3 and MCL1 in spleen samples of control vs KT-333 treated mice, week 2 post-transplantation, 48 hours post drug administration. D) Significant decrease in lineage negative HSPC population along with decrease in intracellular E) total STAT3 F) p-STAT3(S727) and G) p-STAT3(Y705) is observed in the BM cells (p < 0.05) of PDX model collected on week 2, 48 hours post treatment. H) Survival analysis of Ven-res PDX model shows a significant improvement in survival of mice treated with KT-333 as compared to vehicle (Median survival **76** days in vehicle group vs **92** days in KT-333 treated cohort; p = 0.0027).

Next, to evaluate efficacy of KT-333 in Ven-res AML, a murine patient derived xenograft (PDX) model of Ven-res systemic AML was developed ^(35)^. Briefly, NSG mice post irradiation were transplanted with Ven-res AML cell line intravenously 48-hour post-transplantation, the mice were randomized into 2 groups to be treated with vehicle (PBS) or clinical stage STAT3 degrader KT-333 (30mg/kg, IV) once a week. In BM aspirates collected on week 2, 48 hours post treatment, we observed a significant decrease in lineage negative HSPC population along with decrease in intracellular total STAT3 and p-STAT3(S727) in the BM cells (p < 0.05) (Figure 7D-G). Survival analysis shows a significant improvement in survival of mice treated with KT-333 as compared to vehicle (Figure 7H; Median survival **76** days in vehicle group vs **92** days in KT-333 treated cohort; p = 0.0027).

## Discussion

Ven-res is a pervasive and challenging problem to overcome in AML therapy. The median overall survival of AML patients after failure of Ven has been estimated to be 2.4 months ^(36)^. While multiple biological mechanisms of Ven-res have recently been elucidated ^(37)^, most of the approaches targeting these mechanisms have failed clinically or are currently too early in preclinical development ^(38)^.

There are several well characterized mutational drivers of AML and therapeutics targeting these mutations that are in clinical use, the best examples of which are the IDH1/2, FLT3 inhibitors ^(39–43)^. However, these are not very efficacious as single agents and require combination with other therapies for enhanced efficacy ^(44, 45)^. Lesser attention has been paid to clinically targeting downstream mediators of leukemogenesis and particularly transcription factors that are overexpressed in AML due to prior difficulties in developing effective drugs that can bind and downregulate or degrade the transcription factor of interest in leukemic HSPC’s. However, targeting transcription factors represents an attractive and mutation agnostic path forward in AML therapeutics. Degrader technology is now coming of age across several oncologic and non-oncologic indications ^(46–50^). The enhanced specificity towards the protein of interest and selective targeting for degradation make the therapy especially attractive to spur further clinical development and test combinatorial approaches also in preclinical and clinical development in AML.

This paper establishes several new paradigms on STAT3 hyperactivation in AML and shines light on a novel mechanism in therapy-resistant AML. STAT3 hyperactivation is a strong leukemogenic signal that skews HSPCs to a myeloid bias. This signal is further compounded in the setting of therapy resistance in clinical patient datasets where prior exposure to Ven greatly upregulates STAT3 and its phosphorylated forms with a strong association with worsened overall survival. In addition, there is increased translocation of phosphorylated forms of STAT3 into Ven-res mitochondria that correlate with structural and functional mitochondrial defects. A highly specific potent heterobifunctional degrader of STAT3 caused dose dependent and selective degradation of STAT3 as well as p-STAT3(Y705) and p-STAT3(S727) in both parental and Ven-res hematological malignancy cell line. STAT3 degradation not only led to significant induction of apoptosis in cell lines, but also led to reduction in the MCL1 dependent mitochondrial depolarization as observed through BH3 profiling. Additionally, the structural mitochondria defects were restored to levels comparable to healthy subjects upon treatment with the STAT3 degrader. *In-vivo* patient-derived xenograft models of Ven-res AML indicate that effective STAT3 degradation significantly enhances the survival of mice treated with KT-333. STAT3 degradation is a novel and effective strategy with strong mechanistic rationale that can spur further clinical development of STAT3 degraders in Ven-res AML.

In conclusion, even in this era of targeted therapies, only 30% of patients with newly diagnosed acute myeloid leukemia (AML) enjoy long-term survival ^(1)^ while the majority still succumb to their illness. Patients with minimal residual disease (MRD) negative complete remissions also relapse due to chemotherapy-resistant leukemic stem cells (LSCs) ^(51)^. Eradication of residual disease at the LSC level is the ultimate goal of anti-leukemia therapy. We have identified STAT3 to be an important therapeutic target that is overexpressed in LSCs and associated with a poor prognosis and survival. This is even more pronounced in the Ven era where currently there are no available therapies after prior Ven failure. STAT3 and concurrent MCL1 expression is upregulated in Ven-res. A novel STAT3 degrader now shows efficacy in this setting in preclinical models and improves survival in a murine model of Ven-res AML by degrading STAT3 and downregulating MCL1. STAT3 degradation is well poised to enter early phase clinical trials for this relapsed/refractory AML patient population in dire need of newer therapies.

## Materials and Methods

### Patient samples and reagents

MDS, AML and Venetoclax resistant AML patient samples were obtained from an IRB approved biobank at the Albert Einstein College of Medicine. Patient mutation profiles can be accessed from Supplementary Table 2. KTX-201, KTX-105 and KT-333 along with structural controls were provided by Kymera Therapeutics. For in vitro studies, KT-201 and KT-105 were dissolved in DMSO to prepare 20mM stocks. KT-333 was resuspended in PBS for in vivo murine CDX and PDX studies.

### Reverse Phase Protein Array measurement of total STAT3, p-STAT3(Y705) and p-STAT3(S727)

Primary AML patient samples were enriched by ficoll separation and CD3/CD19 depletion, whole cell lysates prepared, then printed onto nitrocellulose slides, stained with highly validated antibodies, and analyzed as previously described ^(52)^. Pearson correlation between total, p-STAT3(Y705) and p-STAT3(S727) forms of STAT3 and the other 426 analysed proteins was performed.

### Cell lines

MOLM13, MOLM16 and SUDHL1 cells were cultured in RPMI (Cytiva) with 10% fetal bovine serum (FBS, Gemini Bio-products) and 1% Penicillin-Streptomycin (Thermo Scientific). Venetoclax resistant cell line MOLM13-VR was generated as previously described ^(53, 54)^ MV411 cells were cultured in IMDM (Fisher Scientific) with 10% fetal bovine serum (FBS) and 1% Penicillin-Streptomycin (P/S). All cells were kept in a 5% CO2 37°C incubator and regularly tested to be free of mycoplasma contamination.

### Mice

NSG Mice were maintained under pathogen-free conditions in a barrier facility in microisolator cages based on a protocol approved by the Institutional Animal Care and Use Committee at Albert Einstein College of Medicine (AECOM).

### Generation of double transgenic STAT3C-vavCre mouse model

To determine the role of STAT3 in myeloid malignancies, a murine model was generated by crossing R26STAT3Cstopfl/fl mice with vavCre transgenic mice. R26STAT3Cstopfl/fl mice were generated and characterized as previously described ^(10)^. Briefly, Bruce4 C57BL/6 embryonic stem cells were transfected with a modified Rosa26 targeting vector that included a 59 floxed stop/Neo cassette and FLAG-tagged Stat3C cDNA, with an frt-flanked IRES-eGFP downstream ^(10)^. Excision of the stop cassette by Cre recombinase leads to expression of a flag-tagged STAT3C protein and concomitant expression of eGFP. R26STAT3Cstopfl/fl and Vav-iCre mice (The Jackson Laboratory) were crossed to obtain desired experimental genotype (R26STAT3Cstopfl/+ VavCre or R26STAT3Cfl/fl VavCre mice). All mice genotypes were determined by PCR and further confirmed by flow cytometry analysis on GFP expression in Hematopoietic stem cells (HSCs). PCR Primers used for mice genotyping are the following: STAT3C forward 5’-GATGCAGTTTGGAAATAACGGTGAA-3’; STAT3C reverse 5’-GAGGTCAGATCCATGTCAAACGT-3’; Rosa26wt forward 5’-TTCCCTCGTGATCTGCAACTC-3’; Rosa26wt reverse 5’-CTTTAAGCCTGCCCAGAAGACT-3’; Vav-iCre forward 5’-TCCTGGGCATTGCCTACAAC-3’; VavCre reverse 5’-CTTCACTCTGATTCTGGCAATTTCG-3’.

Homozygous R26STAT3Cfl/fl VavCre mice were embryonically lethal, therefore heterozygous R26STAT3Cstopfl/+ VavCre mice were utilized for experimental purposes. For most experiments, control animals with a floxed stop cassette but lacking VavCre (R26STAT3Cstopfl/fl) and Vav-iCre control littermates were used. All mice were maintained on a clean C57BL/6 background and housed in specific pathogen-free conditions at the animal facility at Albert Einstein College of Medicine. Animal care was within institutional animal care committee guidelines, and all experiments were performed in accordance with approved protocols for the Albert Einstein College of Medicine Institutional Animal Care and Usage Committee (IACUC).

### Characterization of double transgenic STAT3C-vavCre mice

To determine the role of STAT3 in initiation or progression of myeloid malignancies, heterozygous R26STAT3Cstopfl/+ VavCre mice were utilized for experimental purposes. For most experiments, age-matched control animals with a floxed stop cassette without VavCre (R26STAT3Cstopfl/fl) and Vav-iCre control littermates were used. Bi-weekly CBC analysis and blood smears were performed on heterozygous R26STAT3Cstopfl/+ VavCre mice and control animals for detecting hematological abnormalities that may suggest an underlying malignancy. Mice displaying signs of a hematological malignancy on measuring CBC parameters such as low WBC counts, low platelets etc. in addition to decrease in mice weight, development of ruffled fur etc. were euthanized, followed by necropsy procedure. Bone marrow, Spleen, and Liver tissues were harvested during necropsy and samples were further processed for histopathological analysis or FACS for GFP expression or HSC lineage panel analysis. Spleen organ length and weight were recorded. For FACS HSC lineage panel analysis, the following mouse FACS antibodies were used: CD45-BV650 (Biolegend), Ly6G/C-ef450 (Thermo Fisher Scientific/Invitrogen), CD11b-ef450 (Thermo Fisher Scientific/Invitrogen), CD4-ef450 (Thermo Fisher Scientific/Invitrogen), CD3e-ef450 (Thermo Fisher Scientific/Invitrogen), CD19-ef450 (Thermo Fisher Scientific/Invitrogen), B220-ef450 (Thermo Fisher Scientific/Invitrogen), TER119-ef450 (Thermo Fisher Scientific/Invitrogen), CD34-AF700 (Thermo Fisher Scientific/Invitrogen), CD48-BV605 (BD Biosciences), CD150-PECy7 (Biolegend), c-kit-PE (BD Biosciences), Sca1-APC (Thermo Fisher Scientific/Invitrogen). For dead cell exclusion, cells were stained with Zombie NIR fixable viability kit.

### Serial transplantation STAT3C-Vavcre

Wild-type C57BL/6 CD45.1 (Jackson Laboratories, state) recipient mice (N=10) were irradiated at 550 rad twice at an interval of 4 hours for eradication of innate CD45.1 cells within recipient’s bone marrow. STAT3C-VavCre CD45.2 donor mice (N=5) were euthanized, and bone marrow cells were collected. From one donor animal (STAT3c-VavCre CD45.2), two recipient mice (Wild-type C57BL/6 CD45.1) were transplanted with 3-5 x 10^6^ cells per recipient through tail injection. After cell transplantation, CBC analysis was performed weekly to monitor hematological parameters. After 4-weeks post transplantation, all animals were euthanized and Bone marrow, Spleen, and Liver tissues were harvested during necropsy. Bone marrow cells (pooled femur, tibia, and sternum) were collected by crushing bones in a mortar and pestle using PBS + 2% FBS, and then passed through a 70μM filter for obtaining a single cell suspension. ACK Lysis Buffer (Gibco) was added to each sample for lysis of red blood cells. Samples were washed once with PBS + 2% FBS, and then flow cytometry antibody staining was performed. Mouse FACS antibodies used were the following: CD45.1-PE, CD45.2-APC (Biolegend).

### ExCITE-seq for STAT3C KO murine model

Spleen and bone marrow cells from the femur were collected from two control and two Stat3C KO mice. Each mouse’s cells were individually hash-tagged using the 10X Chromium Next GEM Single Cell 5’ Reagents Kits v2 Dual Index (CG000330 Rev C) as per the manufacturer’s instructions. The cells were then counted using a NucleoCounter-300 automated cell counter with DAPI/AO dye. 2 million cells per sample were resuspended in CITE-seq staining buffer (2% BSA, .01% Tween in PBS) and incubated for 10 minutes with Fc receptor block (TruStain FcX, BioLegend and FcR blocking reagent, Miltenyi) to prevent antibody binding to Fc receptors. The cells were then incubated with hashing and ECCITE-seq surface panel antibodies for 30 minutes at 4°C. The antibodies used for cell hashing and our ECCITE-seq antibody-derived tags (ADT) panel were sourced as TotalSeq-C reagents (BioLegend), and the concentrations and clones used for all conjugated antibodies for staining can be found in supplemental Table x. After staining, the cells were washed three times in PBS containing 2% BSA and .01% Tween, followed by centrifugation (300 ×g for 5 minutes at 4°C) and supernatant aspiration. After the final wash, the cells were resuspended in PBS and filtered through 40-µm cell strainers, combined into two lanes and loaded into the 10x Chromium Single Cell Immune Profiling workflow as per the manufacturer’s instructions.

Post-emulsification, cDNA libraries underwent a serious of amplifications using the 10X Chromium Next GEM Single Cell 5’ Reagents Kits v2 Dual Index (CG000330 Rev C) as per the manufacturer’s instructions. Amplified cDNA’s quality and quantity was assessed on an Agilent BioAnalyzer 210 using a High Sensitivity DNA Kit (Agilent Technologies) and the final libraries on an Agilent TapeStation 420 using High Sensitivity D1000 ScreenTape (Agilent Technologies). The individual libraries were diluted to 2nM and pooled for sequencing. The pools were sequenced with S1 100 Cycle Flow Cell v1.5 run kits (26bp Read1 and 91bp Read2) on the NovaSeq 600 Sequencing System (Illumina), targeting 30,000 and 6,00 reads per cell for the GEX and ADT library, respectively. FASTQ files from the 10x libraries were processed using the count module of Cell Ranger pipeline, version 7.0.0. with Intron mode (10x Genomics) aligned to the mm10 ensemble.

### ExCITE-Seq Analysis

Raw reads were processed using the Cellranger Pipeline (10x Genomics). The expression matrices for individual samples were merged using the Cellranger aggr function. The R package Seurat (v4.3.0) was used to cluster the cells in the merged matrix. Cells with less than 100 genes or more than 1e4 transcripts or 5% of mitochondrial expression were first filtered out as low-quality cells. The NormalizeData function was used to normalize the expression level for each cell with default parameters. The FindVariableFeatures function was used to select variable genes with default parameters. The ScaleData function was used to scale and center the counts in the dataset. Principal component analysis (PCA) was performed on the variable genes. The RunHarmony function from the Harmony package was applied to remove potential batch effect among individual samples. Uniform Manifold Approximation and Projection (UMAP) dimensional reduction was performed using the RunUMAP function. The clusters were obtained using the FindNeighbors and FindClusters functions. The cluster marker genes were found using the FindAllMarkers function. The cell types were annotated by overlapping the cluster markers with the published marker genes. The dot plot was plotted using the DotPlot function. The violin plots were plotted using the VlnPlot function. Differential expression analysis between two group of cells was conducted using the FindMarkers function. Genes with adjusted p value smaller than 0.05 were considered significantly differentially expressed. Enrichr was used for pathway enrichment analysis on the differentially expressed genes.

### Ven resistant Cell derived xenografts

NOD/SCID IL2Rgamma KO NSG mice were initially purchased from Jackson Laboratory and then bred, housed, and handled in the animal facility of Albert Einstein College of Medicine. NSG mice aged 8-10 weeks were sub-lethally irradiated (2.5 Gy) 24 hr prior to injection. 1 x 10^6^ MOLM13 Ven-res cells were administered via tail vein injection. 48-hour post-transplantation, the mice were divided into 2 groups to be treated with STAT3 degrader KT-333 (30mg/kg) or vehicle (PBS). Treatments were done once per week and BM aspirates were collected at week 2 post treatment. BM aspirates were used for flow cytometry based analysis using mice-CD45 FITC (Biolegend) 1:100, and human CD45 PerCPCy5.5 (Biolegend) to observe engraftment and western blot was performed to check for effective STAT3 degradation.

### Ven resistant Patient Derived Xenografts

NOD/SCID IL2Rgamma KO NSG mice were bred, housed, and handled in the animal facility of Albert Einstein College of Medicine. NSG mice aged 8-10 weeks were sub-lethally irradiated (2.5 Gy) 24 hr prior to injection. Mononuclear cells from primary Ven-res AML patients previously reported ^(33)^ were isolated by Ficoll separation. 1 × 10^5^ MNCs were administered via tail vein injection. 3-4 weeks later, BM aspiration were performed and analyzed by flow cytometry for the human cell engraftment utilizing mice-CD45 FITC (Biolegend), and human CD45 PerCPCy5.5 (Biolegend). Mice were considered to be engrafted if they showed 0.1% or higher human derived CD45+ cells. The engrafted mice were randomized for treatment with 30mg/kg KT-333 or vehicle (PBS) once a week. Post 48 hours of treatment, BM aspirations and flow cytometry analysis were performed. Mice were considered severely ill and were euthanized upon reaching a moribund state. Their survival was recorded for the study and endpoint BM and spleen cells were collected and cryopreserved.

### Drug Treatment of Mice

The mice were treated 30mg/kg KT-333 once a week intravenously. The control mice were treated with PBS once a week.

### Bone Marrow Aspirates

For femoral bone marrow aspirations mice will be anesthetized. The animal’s leg was first disinfected with 70% ethanol; a fine needle was then inserted into the femoral bone marrow cavity through the distal condyles. A small volume (up to 10 uL) of bone marrow was aspirated as this has been shown not to compromise the animal’s functionality of the leg or their overall health. Sides (left/right) were alternately used for sequential aspirates. Animals received 5mg/kg Banamine pre-emptive per subcutaneous injection as an analgesic.

### Cell Proliferation Assay

Cell lines and primary samples were incubated at concentrations of 100nM–10 μM of STAT3 degraders - KTX-201 and KTX-105 and controls. Cells were plated in four 96-well plates (1 x 10^4^ cells/well) and treated with KTX-201 and KTX-105 in triplicate for 24 hours, 48 hours, 72 hours. The amount of ATP present was measured using CellTiter-Glo Luminescent Cell Viability Assay (Promega). Cell Titer Glo reagent was added 1:1 before allowing the plate to gently rock 20 minutes at room temperature. Luminescence measured by a Fluostar Omega Microplate reader (BMG Labtech).

### Immunoblotting

Cells were treated with STAT3 degrader (KTX-201, KTX-105, KT-333, Kymera Therapeutics, MA) or vehicle control for 24 hours. Cells were then centrifuged at 350xg for 5 minutes and pellets were collected. Protein lysates were prepared with 1% NP-40 lysis buffer (20 mmol/L Tris-HCl, pH 7.5; 150 mmol/L NaCl; 1 mmol/L EDTA; 150 mmol/l NaCl;1 % NP-40) containing protease inhibitors (Roche) and phosphatase inhibitors cocktail 2 and 3 (Sigma). The cells were lysed for 30 to 45 minutes at 4°C followed by centrifugation at 14000 RPM for 40 minutes at 4°C. Protein quantification was performed using BCA assay. 60 μg of protein was resolved on SDS-PAGE (Biorad; 4–15% Mini-PROTEAN® TGX™ Precast Protein Gels) at constant voltage of 80-100V, followed by transfer to PVDF membrane (EMD-Millipore). Western blot analysis was performed with the following antibodies: STAT3 (Cell Signaling Technology), phospho-Tyr-STAT3 (Cell Signaling Technology), phospho-Ser-STAT3 (Cell Signaling Technology), MCL1 (BD biosciences) and β-actin (Santa-Cruz Biotechnology). Normalisation was done using BioRad Image Lab software.

### Real Time PCR

RNA was isolated from frozen cell pellets using the RNeasy isolation kit (Qiagen) One microgram of total RNA was reverse-transcribed using the iScript cDNA Synthesis Kit (Bio-Rad). qPCR reactions were performed with the SYBR Green PCR Master Mix (Life Technologies) using validated gene-specific primers (Supplemental Table -1). Transcript levels of genes of interest were normalized to a housekeeping gene (GAPDH) and fold change in expression is determined by –ΔΔCt, as previously published ^(9)^.

### Colony Forming Unit assay

0.2-05 million Primary AML PBMNCs and healthy controls were plated in Methocult media (Stem Cell Technologies H4435) in 35×10mm dishes and incubated with and without 1uM KTX-201. The cultures were incubated for 14-18 days. Erythroid and myeloid colonies were then counted, representative colony images were photographed, and the samples were processed for flow cytometry as previously performed ^(9, 55)^

### Caspase 3/7 Assay

MOLM13 Parental and Ven resistant cells (5 × 10^3^ cells/well) were seeded in a 96-well white plate and treated with 100nM KTX-201. Caspase 3/7 activation was measured after 6hr, 24hr and 48 hr by addition of the Caspase-Glo 3/7 reagent according to the manufacturer’s protocol (Promega). Luminescence was detected by a microplate reader (TECAN). Caspase assays were performed in triplicate and the data normalized to vehicle-treated control wells.

### Flow Cytometry

#### Human samples

For human CFU samples, colonies were enumerated after a 14-day incubation period. The CFU cultures in the semisolid media were supplemented with 3 mL of Phosphate-buffered saline containing 2% fetal bovine serum (2% FBS-PBS) and incubated for 1 hour at 37°C in the incubator. After incubation, the culture was thoroughly mixed, transferred into a 15 mL falcon tube, and the cells were pelleted at 350 x g for 10 minutes at 4°C. They were then washed once with phosphate-buffered saline containing 2% fetal bovine serum. Next, the cells were resuspended in 100 µL of Zombie-NIR diluted 1:4000 and incubated in the dark at room temperature for 15 minutes. After the incubation, the cells were washed again with PBS, and the pellet was resuspended in 50 µL of flow antibody cocktail. The mixture was incubated in the dark at room temperature for another 15 minutes. Following another round of washing, the cells were resuspended in PBS. Flow cytometry was then performed on a BD LSRII to acquire the data and analyzed using FlowJo. Erythroid cells were analyzed by staining of unfractionated BMCs with conjugated antibodies against CD71, Glycophorin A, CD11b, CD14, CD233, CD34, CD45 and CD49d (Biolegend).

#### Murine samples

For mouse samples, cells were suspended in sterile fluorescence-activated cell sorting buffer PBS containing 0.5% bovine serum albumin and 2 mM EDTA and stained with indicated surface markers for 30 minutes at 4°C. All flow cytometry data were acquired on a BD LSRII and analyzed using FlowJo. Chimerism in transplantations was assessed by staining PBMCs with conjugated antibodies against CD45.1-APC (Biolegend) and CD45.2-PerCP-Cy5.5 (Biolegend).

### BH3 Profiling

AML cell lines were compared by BH3 profiling under basal conditions by using the plate-based JC-1 BH3 profiling assay ^(56)^. BIM, BID, PUMA, BMF-y, BAD, NOXA, HRK-y, FS1 and MS1 BH3 peptides at indicated concentrations; Puma2A peptide (final concentration of 25 μM); alamethicin (final concentration of 25 μM); CCCP (final concentration of 10 μM) were added to 15μL of JC1-MEB staining solution (20 ug/mL oligomycin, 20 ug/mL digitonin, 2 μM JC-1, 10μM 2-mercaptoethanol in MEB) in a black 384-well plate (Corning, Corning, NY, USA, CLS 3573) using Tecan D300E dispenser. The MEB buffer consisted of 150mM mannitol, 10mM HEPES-KOH pH 7.5, 50mM KCl, 0.02mM EGTA, 0.02mM EDTA, 0.1% BSA and 5mM Succinate.

Single-cell suspensions were washed twice in PBS and resuspended in MEB at 4× their final density (2×10^4^ cells/well). One volume of the 4× cell suspension was added to one volume of the JCI-MEB staining solution. This 2× cell/staining solution was incubated at RT in the dark for 10 min to allow cell permeabilization and dye equilibration. A total of 15 μL of the 2× cell/staining solution mix was then added to each treatment well of the plate. The fluorescence was measured immediately at 590 nm emission 545 nM excitation using the M1000 microplate reader (TECAN) at 30 °C every 15 min for a total of 3 h. Percentage of depolarization was calculated by normalization to the AUC of solvent-only control DMSO (0% depolarization) and the positive control CCCP (100% depolarization), as previously described^(58)^. Bar graph represents the % of mitochondria depolarization of cells detected by JC-1 upon treatment of BH3-derived peptides, n=3. For dynamic BH3-profiling cells were pre-treated with 100nM KTX-201 for 24 hours before BH3 profiling.

### Electron Microscopy

Cell lines and patient derived PBMNCs are centrifuged at 1000 rpm for 10 minutes. This is followed by resuspension of the cells in 1ml IMDM + 2% FBS. The resuspended cells are mixed with 1:1 ratio with fixative and was submitted to EM core at Albert Einstein College of Medicine for imaging. The images were quantified manually and plots were prepared using GraphPad Prism software.

### Immunofluorescence

Cells were seeded on Lab-Tek™ II Chamber Slide (Thermo Fisher Scientific) coated with Retronectin (Clonetech). Samples were fixed by PAF 4% for 10 min at RT, and then permeabilized by Triton 0.25% for 10 min at RT. After blocking with BSA 2% for 30 min at RT, samples were incubated overnight at 4°C with primary Antibodies (dilution 1:50). After washing, samples were incubated with donkey anti-mouse AlexaFluor546 (Invitrogen) and donkey anti-rabbit AlexaFluor488 (Invitrogen) for 1 hr at RT at 1:500 dilutions. After wash, samples were mounted using Vectashield, a mounting medium for fluorescence with DAPI (Vector). Mouse monoclonal anti-TOMM20 (Abnova), rabbit anti-STA3 (Cell Signaling Technology) or rabbit anti-pSTAT3 (Cell Signaling Technology) were used as primary Antibodies. Z-stack were acquired on Leica Stellaris 8 Confocal equipped with 63X oil immersion lens. Stacks were deconvolved using Huygens Essential (SVI). Analysis and representative image renderings were obtained by Imaris 7 (Bitplane).

### Subcellular fractionation

Cell fractionation was performed as previously described ^(57)^. Briefly, 10^9^ cells were harvested in PBS and washed by centrifugation at 500g for 5 min with PBS. The cell pellet was suspended in homogenization buffer [225 mM mannitol, 75 mM sucrose, 30 mM Tris-HCl pH 7.4, 0.1 mM ethylene glycol-bis(β-aminoethylether)-N,N,N′,N′-tetraacetic acid (EGTA), and 1mM PMSF (phenylmethylsulfonyl fluoride)] and gently disrupted by Dounce homogenization. The homogenate was centrifuged twice at 600g for 5 min at 4°C to remove nuclei and unbroken cells, and the resultant supernatant was centrifuged at 7,000g for 10 min at 4°C to pellet crude mitochondria. The supernatant was centrifuged at 20,000g for 30 min at 4°C. Further centrifugation of the supernatant at 100,000g (70-Ti rotor; Beckman) for 90 min at 4°C resulted in the isolation of ER (pellet) and cytosolic fraction (supernatant). To purify mitochondria, the crude mitochondrial fraction was suspended in isolation buffer [250 mM mannitol, 5 mM HEPES pH 7.4, and 0.5 mM EGTA] and subjected to Percoll gradient centrifugation (30% v/v Percoll) in a 10 ml polycarbonate ultracentrifuge tube, at 95,000g (SW41 rotor; Beckman) for 30 min at 4°C. This result in the formation of two rings. The upper one includes mitochondria associated membranes (MAMs) and the lower one isolated mitochondria. Mitochondria-containing ring was then washed by centrifugation at 6,300g for 10 min at 4°C to remove the Percoll and finally suspended in isolation medium.

### Seahorse Mito Stress Test

The Seahorse XF96 Extracellular Flux analyzer (Seahorse Bioscience) was used to measure the oxygen consumption rate (OCR) of MOLM13 cells and MOLM13 Ven res cells according to the manufacturer’s instructions ^(58)^. Briefly, AML cells were cultured with or without 100nM KT-201x for 24 hours. Cells were counted, and 7.5 × 10^5^ cells were added to each well for the extracellular flux assays. Three technical replicates for each condition were plated.

### Statistical analyses

Data visualization was done using GraphPad Prism 10 software. For the comparison between two experimental groups, an unpaired t-test was used. For more than two groups, ANOVA test was used with Tukey’s multiple comparisons test. In all graphs, error bars indicate mean ± SEM. A P value less than 0.05 was considered significant (*), with 0.01 (**), 0.001 (***), and 0.0001 (****) representing higher levels of significance. Non-normally distributed data was analyzed using Mann-Whitney’s test. Overall survival was measured and analysed by Kaplan-Meier plotting using GraphPad Prism software. Immunoblot image quantitation was based on at least three independent biological replicates.

## Acknowledgements

We thank the MDS/AML patients for their contributions. We thank Swathi of the AECOM Stem Cell Isolation and Xenotransplantation Facility (funded through New York Stem Cell Science grant no. C029154), Leslie Gunther Cummins, Frank Macaluso and Joseph Churaman from Analytical Imaging Facility (funded through NCI cancer centre support grant P30CA013330 and shared instrument grant JEOL 1400Plus TEM: SIG #1S10OD016214-01A1), Dr. Xue-Liang Du from Stable Isotope and Metabolomics Core and Jinhang Zhang, M. Liu and A. Feng from the AECOM Flow Cytometry Core Facility for assistance with flow cytometry. We also acknowledge the availability of Leica Stellaris 8 Confocal Microscope (1S10OD034397-01) in the shared facility of Albert Einstein College of Medicine.

## Supplementary Tables

**Supplementary Tables 1:**
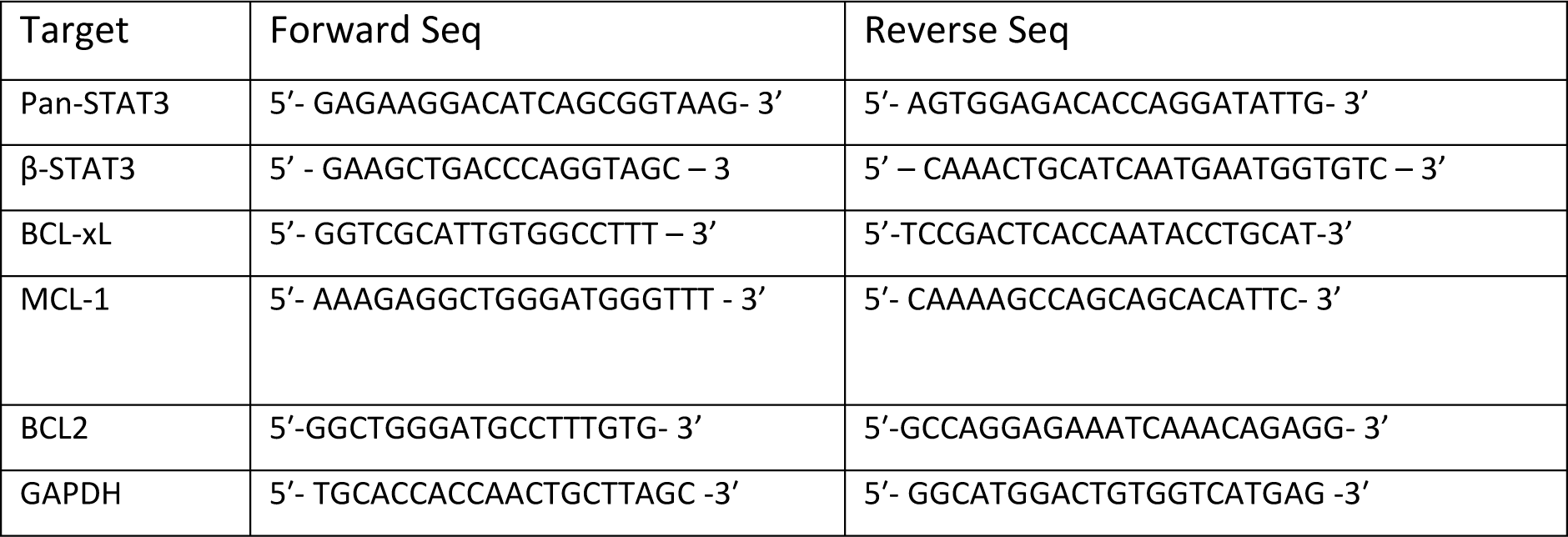
RT-PCR Primer sequences.

**Supplementary Tables 2:** Patient mutation profile.

2. Patient mutation profile
| Patient Name | Mutation Profile |
| --- | --- |
| Sample C | JAK2V617F mutation (33.3% VAF), FISH shows trisomy 8 and complex cytogenetics. |
| Sample 427 | JAK2 - VAF: 94%; IDH1 - VAF: 47%; NPM1 - VAF: 44% |
| Sample 101 | NPM1 - VAF: 20% |
| Sample 417 | TET2 – VAF: 86% ; GATA2 – VAF: 44% ; CEBPA – 2 mutations – VAF: 29% (pathogenic); 32% (likely pathogenic) |
| Sample 300 | NPM1 – VAF: 49% ; TET2 – VAF: 40% |
| Sample 129 | FLT3 Internal Tandem Duplication with signal ratio of >1 |

## Supplementary Figures

Supplementary Figure 1. A, B) Phospho-proteomic analysis on AML patients treated with Ven shows significant reduction in remission duration (RemDur) in patients with high expression of p-STAT3(Y705), p-STAT3(S727) respectively.

Supplementary Figure 2. A) Western blot showing no effect on the expression of STAT5 in MOLM13 parental cells when treated with STAT3 degraders (D1: KTX-201, D2: KTX-105) and their structural controls (C1, C2) at 0.1, 1 and 10μM doses for 24 hours. B) Western blot showing no effect on the expression of STAT5 in MOLM13 Ven-res cells when treated with STAT3 degraders (D1: KTX-201, D2: KTX-105) and their structural controls (C1, C2) at 0.1, 1 and 10μM doses for 24 hours.

Supplementary Figure 3. A) FACS post CFU assay using AML patient PBMNC shows increased differentiation of early erythroid markers CD71 vs GlyA on treatment with KTX-201. B) FACS post CFU assay using Ven-res AML patient PBMNC shows increased differentiation of early erythroid markers CD71 vs GlyA from 7.7% to 12.6% on treatment with KTX-201. C) No differentiation in erythroid markers CD71 and GlyA was observed in PBMNC sample from healthy subject on treatment with KTX-201.

## References

1. Shastri, A., et al., Stem and progenitor cell alterations in myelodysplastic syndromes. Blood, 2017. 129(12): 1586–1594

2. DiNardo, C., et al., Venetoclax combined with FLAG-IDA induction and consolidation in newly diagnosed acute myeloid leukemia. Am J Hematol, 2022.97(8):1035–1043

3. Mantzaris, I., et al., Venetoclax Combined with “7+3” Induction Chemotherapy Induces High Rates of MRD-Negative Remission in Newly Diagnosed AML Patients Fit for Intensive Chemotherapy Irrespective of Age. Blood, 2023. 142 (Supplement 1): 4257

4. Kanna R, Choudhary G, Ramachandra N, Steidl U, Verma A, Shastri A. STAT3 inhibition as a therapeutic strategy for leukemia. Leuk Lymphoma, 2018;59(9):2068–2074

5. Munoz, J., et al., STAT3 inhibitors: finding a home in lymphoma and leukemia. Oncologist, 2014. 19(5): 536–44

6. Xiong, A., et al., Transcription Factor STAT3 as a Novel Molecular Target for Cancer Prevention. Cancers, 2014. 6(2): 926–957

7. Zhang, Q., et. al,. Mitochondrial localized Stat3 promotes breast cancer growth via phosphorylation of serine 727. J Biol Chem. 2013. 288(43):31280–8

8. Beebe JD., et. al., Two decades of research in discovery of anticancer drugs targeting STAT3, how close are we? Pharmacol Ther. 2018. 191:74–91.

9. Shastri, A., et al., Antisense STAT3 inhibitor decreases viability of myelodysplastic and leukemic stem cells. J Clin Invest, 2018. 128(12): 5479–5488

10. Fogli, L. K., et al., T cell-derived IL-17 mediates epithelial changes in the airway and drives pulmonary neutrophilia. J Immunol. 2013. 191(6): 10.4049

11. Rivera, B., et. al., A Transgenic Murine Model Expressing Hyperactive STAT3 Recapitulates the Features of MDS/AML. Blood. 2021. 138 (Supplement 1): 3308

12. Stahl, M., et. al., Clinical and molecular predictors of response and survival following venetoclax therapy in relapsed/refractory AML. Blood Adv. 2021. 5(5):1552–64.

13. Zhang, H., et. al., Integrated analysis of patient samples identifies biomarkers for venetoclax efficacy and combination strategies in acute myeloid leukemia. Nat Cancer. 2020. 1(8):826–39.

14. Zhang, Q., et al., Activation of RAS/MAPK pathway confers MCL-1 mediated acquired resistance to BCL-2 inhibitor venetoclax in acute myeloid leukemia. Signal Transduct Target Ther. 2022. 7(1):51

15. Nishida, Y., et al., Enhanced TP53 reactivation disrupts MYC transcriptional program and overcomes venetoclax resistance in acute myeloid leukemias. Sci Adv. 2023. 9(48):1436

16. Csibi, F., et al., Small Molecule-Induced, Selective STAT3 Degradation Leads to Anti-Tumor Activity in STAT3-Dependent Heme Malignancies. Blood. 2019. 134 (Supplement 1): 3803

17. Sharma, K., et al., E3 pairing and structural mechanisms underlying anti-tumor activity of clinical STAT3 degrader KT-333. Cancer Res. 2024. 84 (Supplement 7): LB037

18. Ebert, BL., et al., An erythroid differentiation signature predicts response to lenalidomide in myelodysplastic syndrome. PLoS Med. 2008. 5(2): e35

19. Sperling AS, Gibson CJ, Ebert BL. The genetics of myelodysplastic syndrome: from clonal haematopoiesis to secondary leukaemia. Nat Rev Cancer. 2017. 17(1):5–19

20. Chen, X., et al., Targeting Mitochondrial Structure Sensitizes Acute Myeloid Leukemia to Venetoclax Treatment. Cancer Discov. 2019. 9(7):890–909.

21. Perciavalle, RM., et al., Anti-apoptotic MCL-1 localizes to the mitochondrial matrix and couples mitochondrial fusion to respiration. Nat Cell Biol. 2012. 14(6):575–83.

22. Wegrzyn J., et al., Function of mitochondrial Stat3 in cellular respiration. Science. 2009. 323(5915):793–7.

23. Meier, J A., et al., Toward a new STATe: the role of STATs in mitochondrial function. Semin Immunol. 2014. 26(1):20–8.

24. Szczepanek, K., et. al., Mitochondrial-targeted Signal transducer and activator of transcription 3 (STAT3) protects against ischemia-induced changes in the electron transport chain and the generation of reactive oxygen species. J Biol Chem. 2011. 286(34):29610–20.

25. Zhang, Q., et al., Mitochondrial localized Stat3 promotes breast cancer growth via phosphorylation of serine 727. J Biol Chem. 2013. 288(43):31280–8.

26. Capron, C., et al., Viability and stress protection of chronic lymphoid leukemia cells involves overactivation of mitochondrial phosphoSTAT3Ser727. Cell Death Dis. 2014.5(10):e1451.

27. Gough, D J., et. al., Mitochondrial STAT3 supports Ras-dependent oncogenic transformation. Science. 2009. 324(5935):1713–6.

28. Szczepanek, K., et. al., Mitochondrial-targeted Signal transducer and activator of transcription 3 (STAT3) protects against ischemia-induced changes in the electron transport chain and the generation of reactive oxygen species. J Biol Chem. 2011. 286(34):29610–20.

29. Rincon, M., et. al., A New Perspective: Mitochondrial Stat3 as a Regulator for Lymphocyte Function. Int J Mol Sci. 2018. 19(6).

30. Zhou, L., et. al., Mitochondrial localized STAT3 is involved in NGF induced neurite outgrowth. PLoS One. 2011. 6(6):e21680.

31. Cogliati, S., et. al., Mitochondrial Cristae: Where Beauty Meets Functionality. Trends Biochem Sci. 2016. 41(3):261–73.

32. Cogliati, S., et al. Mitochondrial cristae shape determines respiratory chain supercomplexes assembly and respiratory efficiency. Cell. 2013. 155(1):160–71.

33. Guieze R., et al., Mitochondrial Reprogramming Underlies Resistance to BCL-2 Inhibition in Lymphoid Malignancies. Cancer Cell. 2019. 36(4):369–84 e13.

34. Marchetti P., et al., Mitochondrial spare respiratory capacity: Mechanisms, regulation, and significance in non-transformed and cancer cells. FASEB J. 2020. 34(10):13106–24.

35. Verma, D., et al., Glutaminase inhibition in combination with azacytidine in myelodysplastic syndromes: Clinical efficacy and correlative analyses, Nature Medicine, 2024.

36. Maity, A., et al., Outcomes of relapsed or refractory acute myeloid leukemia after frontline hypomethylating agent and venetoclax regimens. Hematologica. 2021. 106 (3)

37. Desai, S., et al., Mechanisms of resistance to hypomethylating agents and BCL-2 inhibitors. Best Practice & Research Clinical Haematology. 2023. 36 (4)

38. DiNardo, C., et al., Molecular patterns of response and treatment failure after frontline venetoclax combinations in older patients with AML. Blood. 2020. 135(11): 791–803

39. Issa, G., et al., Acute myeloid leukemia with IDH1 and IDH2 mutations: 2021 treatment algorithm. Blood Cancer Journal. 2021. 11 (107)

40. DiNardo, C., et al., Durable Remissions with Ivosidenib in IDH1-Mutated Relapsed or Refractory AML. N Engl J Med. 2018. 378(25):2386–2398

41. Erba, H., et al., Quizartinib plus chemotherapy in newly diagnosed patients with FLT3-internal-tandem-duplication-positive acute myeloid leukaemia (QuANTUM-First): a randomised, double-blind, placebo-controlled, phase 3 trial. Lancet. 2023. 401(10388):1571–1583

42. Perl, A., et al., Gilteritinib or Chemotherapy for Relapsed or Refractory FLT3-Mutated AML. N Engl J Med. 2019. 381:1728–1740

43. Stone R., et al., Midostaurin plus Chemotherapy for Acute Myeloid Leukemia with a FLT3 Mutation. N Engl J Med. 2017. 377:454–464

44. Montesinos, P., et al., Ivosidenib and Azacitidine in IDH1-Mutated Acute Myeloid Leukemia. N Engl J Med. 2022. 386:1519–1531

45. Kindler, T., et al., FLT3 as a therapeutic target in AML: still challenging after all these years. Blood. 2010. 116 (24): 5089–5102

46. Bidard, F., et al., Elacestrant (oral selective estrogen receptor degrader) Versus Standard Endocrine Therapy for Estrogen Receptor–Positive, Human Epidermal Growth Factor Receptor 2–Negative Advanced Breast Cancer: Results From the Randomized Phase III EMERALD Trial. Journal of Clinical Oncology. 2022. 40(28)

47. Gao, X., et al., Phase 1/2 study of ARV-110, an androgen receptor (AR) PROTAC degrader, in metastatic castration-resistant prostate cancer (mCRPC). Journal of Clinical Oncology. 2022. 40(6)

48. Campone, M., VERITAC-2: A global, randomized phase 3 study of ARV-471, a proteolysis targeting chimera (PROTAC) estrogen receptor (ER) degrader, vs fulvestrant in ER+/human epidermal growth factor receptor 2 (HER2)-advanced breast cancer. Journal of Clinical Oncology. 2023. 41(16)

49. Khan, S., et al., A selective BCL-XL PROTAC degrader achieves safe and potent antitumor activity. Nat. Med. 2019. 25, 1938–1947

51. Lovisa, F., et al., Pre-and post-transplant minimal residual disease predicts relapse occurrence in children with acute lymphoblastic leukaemia. Br J Haematol. 2018. 180(5):680–693

52. de Camargo Magalhães, E S., et. al., Proteomics for optimizing therapy in acute myeloid leukemia: venetoclax plus hypomethylating agents versus conventional chemotherapy. Leukemia. 2024. 38(5):1046–56.

53. Choudhary, G S., et. al., MCL-1 and BCL-xL-dependent resistance to the BCL-2 inhibitor ABT-199 can be overcome by preventing PI3K/AKT/mTOR activation in lymphoid malignancies. Cell Death Dis. 2015.6(1):e1593.

54. Chakraborty, S., et. al., A STAT3 Degrader Demonstrates Pre-Clinical Efficacy in Venetoclax Resistant Acute Myeloid Leukemia. Blood. 2023. 142 (Supplement 1): 2787.

55. Schinke C, Giricz O, Li W, Shastri A, Gordon S, Barreyro L, et al. IL8-CXCR2 pathway inhibition as a therapeutic strategy against MDS and AML stem cells. Blood. 2015.125(20):3144–52.

56. Ryan, J., et. al., BH3 profiling in whole cells by fluorimeter or FACS. Methods. 2013. 61(2):156-64.

57. Wieckowski, M R., et. al., Isolation of mitochondria-associated membranes and mitochondria from animal tissues and cells. Nat Protoc. 2009. 4(11):1582–90.

58. Vangapandu, H V., et. al., B-cell Receptor Signaling Regulates Metabolism in Chronic Lymphocytic Leukemia. Mol Cancer Res. 2017. 15(12):1692–703.

