## Supplementary figures and images for "A STAT3 Degrader Demonstrates Pre-clinical Efficacy in Venetoclax resistant Acute Myeloid Leukemia"

### Supplementary figures

#### Supplementary Figure 1

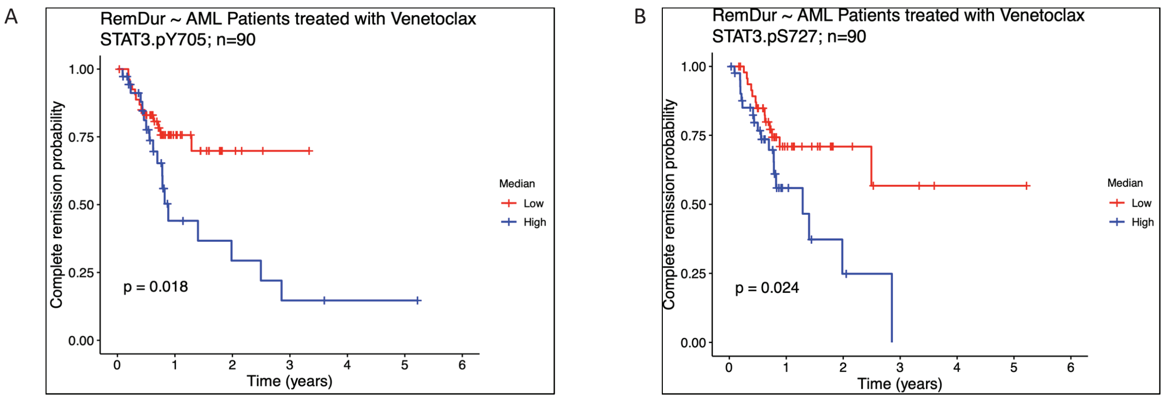


#### Supplementary Figure 2

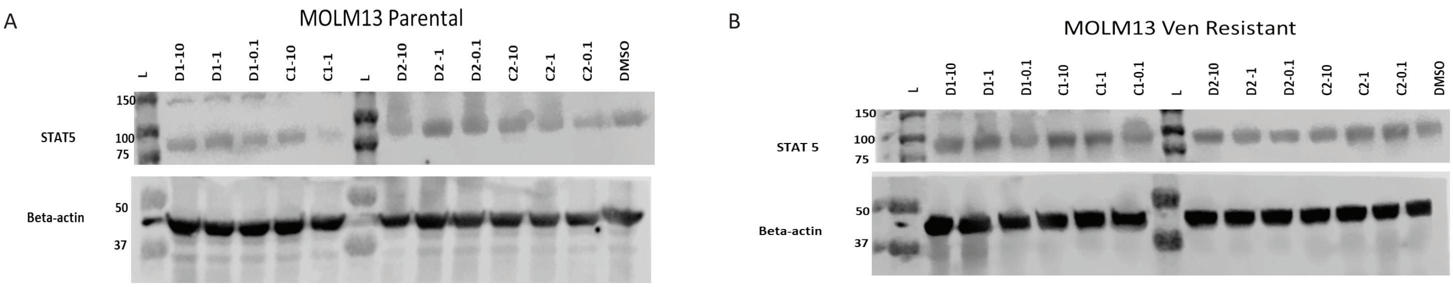


#### Supplementary Figure 3

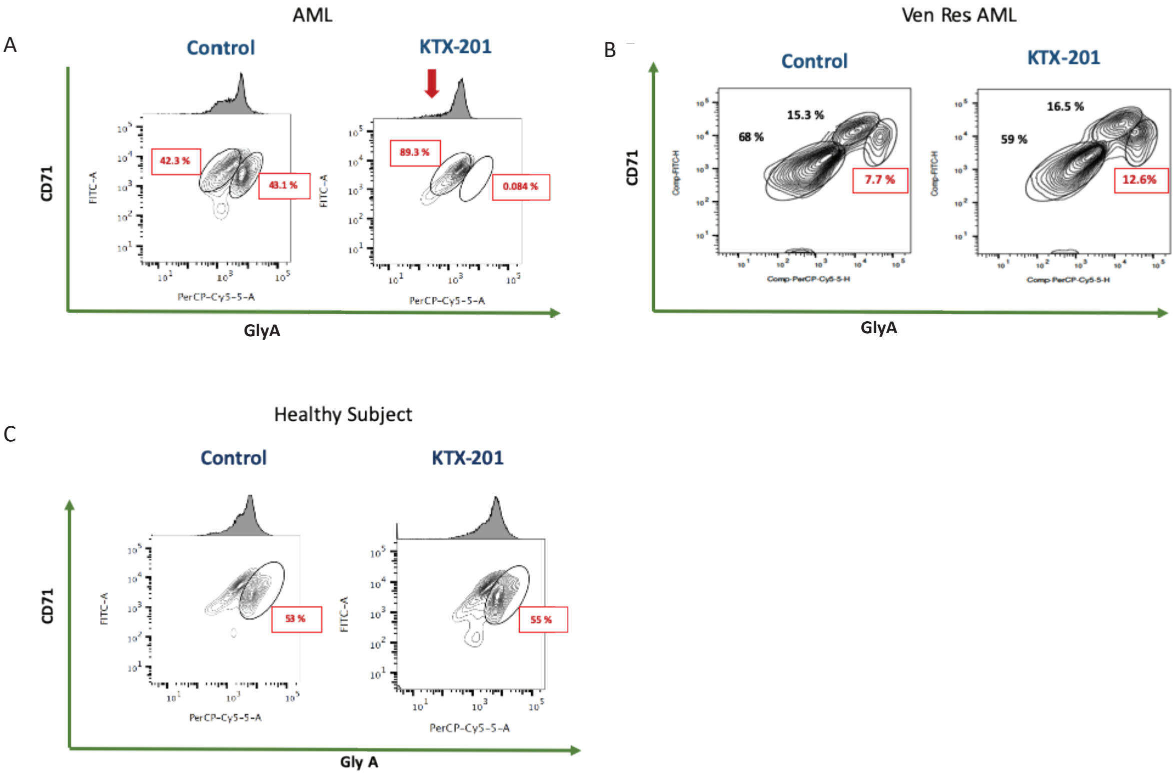
